# Parallel hierarchical encoding of linguistic representations in the human auditory cortex and recurrent automatic speech recognition systems

**DOI:** 10.1101/2025.01.30.635775

**Authors:** Menoua Keshishian, Gavin Mischler, Samuel Thomas, Brian Kingsbury, Stephan Bickel, Ashesh D. Mehta, Nima Mesgarani

## Abstract

Transforming continuous acoustic speech signals into discrete linguistic meaning is a remarkable computational feat accomplished by both the human brain and modern artificial intelligence. A key scientific question is whether these biological and artificial systems, despite their different architectures, converge on similar strategies to solve this challenge. While ASR systems now achieve human-level performance, research on their parallels with the brain has been limited by biologically implausible, non-causal models and comparisons that stop at predicting brain activity without detailing the alignment of the underlying representations. Furthermore, studies using text-based models overlook the crucial acoustic stages of speech processing. Here, using high-resolution intracranial recordings and a causal, recurrent ASR model, we bridge these gaps by uncovering a striking correspondence between the brain’s processing hierarchy and the model’s internal representations. Specifically, we demonstrate a deep alignment in their algorithmic approach: neural activity in distinct cortical regions maps topographically to corresponding model layers, and critically, the representational content at each stage follows a parallel progression from acoustic to phonetic, lexical, and semantic information. This work thus moves beyond demonstrating simple model-brain alignment to specifying the shared underlying representations at each stage of processing, providing direct evidence that both systems converge on a similar computational strategy for transforming sound into meaning.

## Introduction

Understanding how the human brain processes and encodes linguistic information is a fundamental challenge in neuroscience and artificial intelligence. The human auditory cortex is capable of extracting meaning and structure from spoken language with remarkable efficiency. In parallel, automatic speech recognition (ASR) systems have achieved near-human accuracy in recognizing and transcribing speech (*1*). However, it remains unclear how closely the internal computation and representation of these ASR systems mirror those of the human brain, leaving a substantial gap in our understanding of speech processing in humans and machines. Are both systems converging toward similar strategies, or are they reaching different solutions independently as they optimize for performance?

Neuroimaging studies of speech processing in the brain have revealed an emergent encoding of linguistic hierarchies, progressing from primary to nonprimary areas of the auditory cortex (*2–4*). These studies show that the brain distributes, yet jointly encodes, various linguistic features, including phonemes, phonotactics, lexical-phonological, and lexical-semantic information. However, parallels between these hierarchical patterns in the brain and those found in ASR algorithms have not been directly established. Several studies have examined how end-to-end ASR systems represent linguistic information (*5*) and explored similarities between these systems and the brain (*6*). Additional research has also investigated the representational similarities between large language models (LLMs) and the brain (*7–11*). While these studies have provided valuable insights, they have some limitations. Many lacked the temporal precision necessary for speech processing, as they relied on functional MRI (fMRI) (*7*, *10*) which is slow in capturing the neural dynamics. Moreover, by comparing the brain to text-based LLMs, these studies inherently overlooked the acoustic stages of speech processing even though the subjects listened to the stimuli (*7–11*). Other studies used models that were biologically implausible, such as transformers (*6–11*) or non-causal architectures (*6*). In addition, most studies did not explicitly analyze linguistic representation in the models (*7*, *9–11*), or used only a narrow set of features (*6*, *8*). Therefore, even though it has been shown that speech processing models and the brain use increasingly similar representations as revealed by the predictability of neural responses from these models, the precise nature of this convergent similarity remains unclear, replacing one black box with another one without explaining the full picture of speech understanding (*12*).

This study addresses several gaps in the current understanding of speech processing in both biological and artificial systems. First, we expand on previous studies by investigating a broad range of linguistic features, providing a comprehensive analysis that includes phonetic, lexical, semantic, and contextual representations. Second, we use a biologically plausible recurrent neural network transducer (RNN-T) (*13*) model that processes speech in a causal and incremental manner, aligning more closely with how the human brain processes speech in real time. By incorporating high-resolution intracranial electroencephalography (iEEG) data from participants listening to continuous speech, we establish a direct comparison between neural activations and the internal states of the ASR model.

Through a detailed node-level analysis, we compare specific neural sites in the brain with individual nodes in the ASR model to directly assess representational alignment. Additionally, a layer-level analysis provides a more comprehensive view of the ASR model’s internal hierarchical representations, allowing us to examine how the structure of these representations parallels the cortical encoding of speech in humans. We not only directly compare the brain with the model, but we also explicitly test the linguistic features encoded within both the brain and model, moving beyond simply mapping black-box representations onto neural responses. This dual approach enables us to uncover both fine-grained and hierarchical similarities in how speech is processed by biological and artificial systems, shedding light on shared mechanisms of linguistic encoding while also revealing key divergences.

## Results

We recorded intracranial electroencephalography (iEEG) data from fifteen human participants implanted with subdural (electrocorticography; ECoG) and depth (stereotactic EEG; sEEG) electrodes. The participants listened to 30 minutes of continuous speech spoken by four speakers. To ensure that the subjects were engaged in the task, we paused the audio at random intervals and asked the subjects to report the last sentence before the pause. All subjects were attentive and could correctly repeat the speech utterances with 100% accuracy. We extracted the envelope of the high-gamma frequency band (70-150 Hz), which has been shown to correlate with neural firing in the proximity of the recording electrode (*14*, *15*), as the neural response measure of the recorded signals.

We restricted our analyses to electrode sites in the auditory cortex (AC; 𝑁 = 335) and the inferior frontal gyrus (IFG; 𝑁 = 191). Figure 1a shows the general location of the IFG and the subregions of the auditory cortex on the FreeSurfer average brain (*16*). We further limited our analysis to sites that were determined to be speech-responsive (𝑁 = 291/526, determined by a t-test between responses during speech and in silence), including 62 ECoG electrodes and 229 sEEG electrodes. We labeled the neural sites in both hemispheres based on their anatomical region to enable population tests: posteromedial Heschl’s gyrus (pmHG; 𝑁_𝐿_ = 12/15, 𝑁_𝑅_ = 16/19), anterolateral Heschl’s gyrus (alHG; 𝑁_𝐿_ = 34/36, 𝑁_𝑅_ = 32/34), planum temporale (PT; 𝑁_𝐿_ = 9/12, 𝑁_𝑅_ = 18/20), middle superior temporal gyrus (mSTG; 𝑁_𝐿_ = 44/53, 𝑁_𝑅_ = 17/18), posterior superior temporal gyrus (pSTG; 𝑁_𝐿_ = 27/32, 𝑁_𝑅_ = 21/23), anterior superior temporal gyrus (aSTG; 𝑁_𝐿_ = 19/60, 𝑁_𝑅_ = 0/13), and inferior frontal gyrus (IFG; 𝑁_𝐿_ = 32/119, 𝑁_𝑅_ = 10/72). The electrode locations and their responsiveness are plotted in Figure 1b on the average FreeSurfer brain, where the color indicates whether an electrode is speech-responsive.

**Figure 1.**
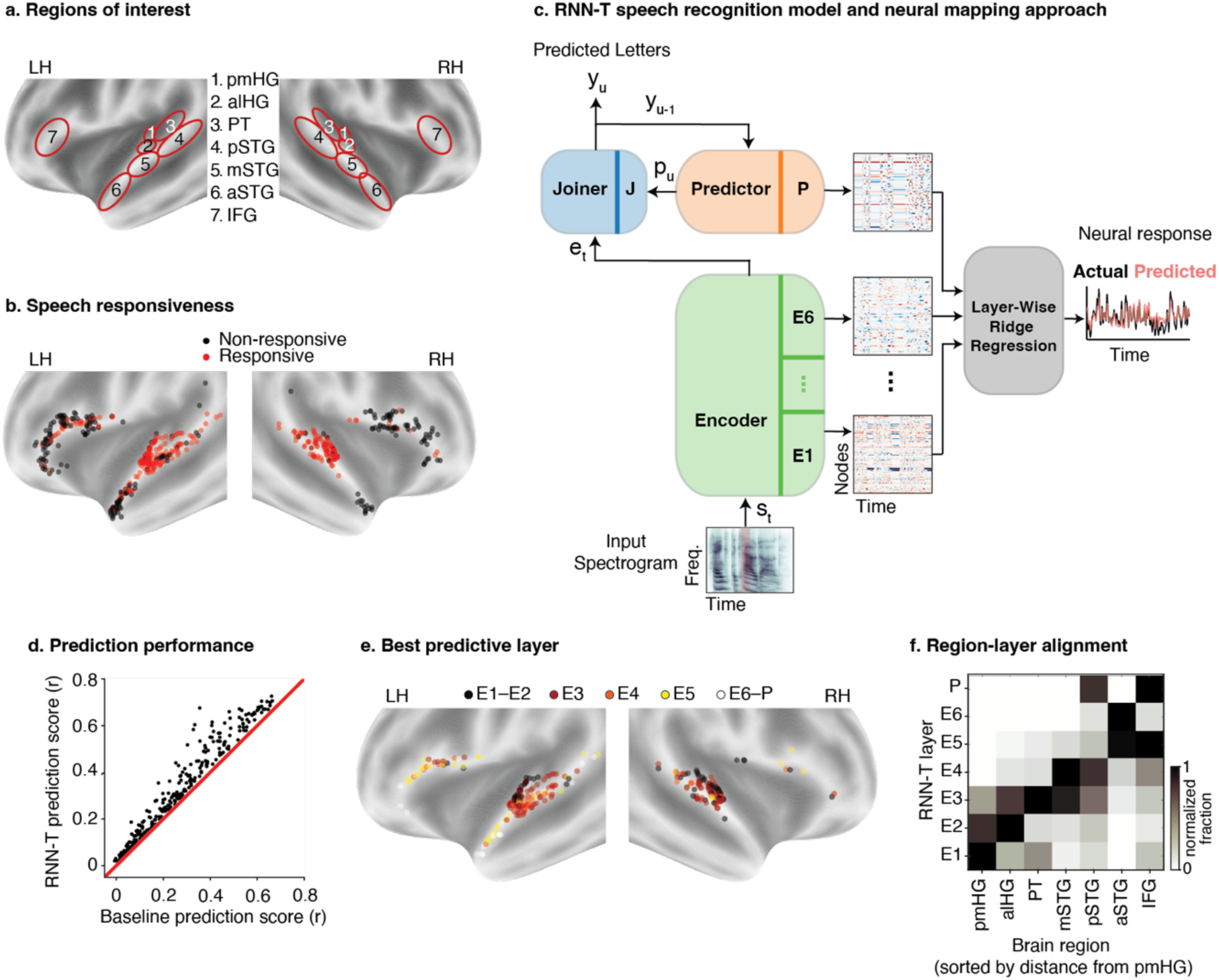
Stages of speech processing in the brain based on ASR modelling. (a) General location of the inferior frontal gyrus (IFG) and subregions of the auditory cortex (AC). (b) Electrode locations within the regions of interest. Colours indicate whether an electrode showed significantly higher activation during speech stimuli compared to pre-trial silence. (c) The RNN-T model consists of a bottom-up Encoder (green) which processes the input spectrogram to produce acoustic representations (layers E1–E6), and a top-down Predictor (orange) which processes the label history to produce linguistic representations (layer P). A Joiner network combines these to predict output labels. To map model representations to neural activity, we fit a layer-wise Ridge regression model to predict the neural response at each electrode from the activations of each specific model layer. (d) Comparison of the cross-validated prediction score using the best ASR layer versus a baseline spectrogram predictor for all speech-responsive electrodes (N=291). (e) Electrodes are coloured according to the ASR model layer (E1–E6, P) that predicts their activity with the highest accuracy. (f) The normalized fraction of electrodes within each anatomical subregion that are best predicted by each layer of the ASR model. Each row is normalized by its maximum value to highlight the subregion where a given layer is most strongly represented. Regions are sorted by their average distance from pmHG.

### Stages of speech processing in the brain based on ASR modeling

We use an RNN-Transducer (RNN-T) (*13*), a recurrent neural network trained to predict letters (graphemes) from the speech spectrogram, to model the speech recognition process in the brain. RNN-T models take spectro-temporal speech signals as input through a causal mechanism, and have been widely used for speech recognition, including for recent state-of-the-art systems in certain domains (*17–22*). Unlike transformer-based architectures, RNN-T models can be causal and recurrent, making them an ideal candidate model to compare with the full hierarchy of speech processing in the brain. The model consists of two branches: the encoder and the predictor (Figure 1c. The encoder branch, consisting of 6 uni-directional LSTM layers (E1–E6), acts as a bottom-up encoder by producing acoustic embeddings from the input speech spectrogram. The predictor branch, consisting of 1 uni-directional LSTM layer (P), acts as an internal language model in that it is conditioned on previous non-blank symbols produced by the model, performing the role of top-down cortical feedback on language likelihood in real time. It stores an internal representation of the prior predictions of the model that together with the encoder’s output are combined by a shallow joint network and used to make the next prediction. We computed layer activations for all layers of these two branches in response to the same 30- minute stimulus set that the human participants listened to.

To examine how the speech processing pathways of the brain and the model map to each other, we find the best matching model layer for each electrode site. First, for each electrode-layer pair, we fit a single-lag regression model that predicts the electrode activity from the layer activations in response to the same stimuli (reduced to top 256 principal components). By using a single-lag framework, we conceptualize each ASR layer as a candidate model for the computation occurring at a given neural site. This approach treats the neural activity as a subspace of the layer’s representational space at a single point in time, which is consistent with the causal, step-by-step processing of an RNN where each layer receives a single time-snapshot from the layer preceding it. The model is defined as follows:

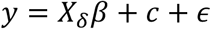

where 𝑦 is the 𝑇 × 1 response vector for an electrode site, 𝑋_#_ is the 𝑇 × 𝐻 matrix of layer activations shifted by a constant time lag 𝛿, 𝛽 is the 𝐻 × 1 vector of regression coefficients, 𝑐 is a constant bias term and 𝜀 is the regression error. We also define a baseline 256-dimensional predictor which is obtained from a 128-dimensional log-Mel-spectrogram stacked on top of a 1- sample delayed version of itself to capture local temporal spectral changes. For each electrode, we also fit a single-lag linear regression model that predicts the neural activity from the baseline predictor, allowing for a fixed time lag 𝛿. Figure 1d shows the improvement of cross-validated prediction scores for each electrode when predicting from the best-performing RNN-T layer compared to the baseline spectrogram predictor. The RNN-T model significantly outperforms the baseline spectrogram model (average correlation improvement = 0.053, 𝑝 ≪ 1𝑒 − 4, paired t-test).

To contextualize this performance, we compared the task-optimized RNN-T against two alternative architectures: a Conformer-based ASR model (*23*) trained on the same speech recognition task, and a CNN-based encoding model trained explicitly to predict the neural activity (*24*, *25*). The Conformer ASR model was trained using a recent training recipe (*26*) on the same training data as the RNN-T and achieved a word error rate of 6.9% on the stimuli used in this study (compared to 4.9% by the RNN-T). We find that the RNN-T achieves very similar prediction scores to the Conformer ASR model (Wilcoxon ranksum test, p>0.05). Additionally, despite never being exposed to neural data during its training, the RNN-T achieves prediction scores on par with the specialized CNN encoding model (Supplementary Fig. S1).

We next associate each electrode with the layer of the RNN-T model that predicts it with the highest accuracy. The colors in Figure 1e indicate the corresponding layer of the model for each neural site in the AC and IFG; and Figure 1f shows the normalized fraction of electrodes in each brain region best predicted by each layer of the model (values indicate fraction of neural sites in brain region, divided by the maximum fraction within layer/row). The model layers map to the cortex such that as we move deeper in the model, in the cortex we move from the primary auditory cortex (pmHG) laterally to PT and mSTG, and from there to pSTG, aSTG, and IFG. In Figure 1f, the brain regions are sorted from nearest to farthest from primary auditory cortex. The corresponding model depth (layer number) of an electrode is also correlated with various other metrics associated with being downstream in speech processing – relative prediction improvement from ASR over baseline (Spearman’s 𝑟 = 0.582, 𝑝 ≪ 1×10^−4^), neural response latency (𝑟 = 0.373, 𝑝 ≪ 1×10^−4^), neural site’s distance from the primary auditory cortex (center of pmHG chosen as reference for primary AC; 𝑟 = 0.613, 𝑝 ≪ 1×10^−4^). Taken together, these results confirm a robust linear alignment between the model’s computational depth and the brain’s biological hierarchy, defined by both anatomical distance from the primary auditory cortex and the temporal latency of neural responses The brain results, however, show a difference in encoding between hemispheres, where neural sites corresponding to deeper layers of the ASR model are predominantly found in the left hemisphere. We can see this by a statistical comparison between the distribution of associated model depth of electrodes (one-sided Wilcoxon rank-sum test, 𝑈 = 5.61, 𝑝 = 1.0 × 10^−8^, 177 speech-responsive sites in left and 114 in right). We also observe a difference by anatomical region. Specifically, although the electrode coverage is not uniform across regions in both hemispheres, a pattern emerges whereby lateralization increases as we move further away from the primary AC – pmHG (𝑈 = −0.139, 𝑝 = 0.56, 12 left, 16 right), alHG (𝑈 = −1.14, 𝑝 = 0.87, 34 left, 32 right), PT (𝑈 = 1.90, 𝑝 = 0.029, 9 left, 18 right), mSTG (𝑈 = 2.02, 𝑝 = 0.022, 44 left, 17 right), pSTG (𝑈 = 2.11 𝑝 = 0.017, 27 left, 21 right), aSTG (no speech-responsive sites in the right hemisphere), IFG (𝑈 = 2.27, 𝑝 = 0.011, 32 left, 10 right). These results are in line with previous studies showing left-lateralization of higher-order regions in linguistic processing (*27*, *28*). Lower-level acoustic processing (according to ASR) is similar between hemispheres, while higher-level processing is biased towards the left hemisphere.

### Similar node-level linguistic encoding across the brain and ASR

We used a regression-based method with temporal receptive fields (TRFs) (*29*) to measure the degree of encoding of different levels of linguistic information in individual nodes of the ASR model and individual neural sites (Figure 2a). This method stands in contrast to the previous single-lag model, but in this case, a window of delays is more appropriate than a single-lag model for capturing the joint encoding of multiple features that may have varying latencies at a single neural site. In this method, we first predicted the neural activity of a site from a broad set of acoustic and linguistic features, including pitch, phonemes, phonotactics, lexical entropy and surprisal, word frequency, and semantic neighbourhood density (see Methods for definitions) (*30*). Next, we determined the contribution of each feature in predicting the response by replacing it with a control variable and interpret the drop in predictive power as the contribution of that feature to the prediction (*30*). We repeated the replacement process for each linguistic feature 100 times and measured the t-statistic between actual and control features to determine the significance of the feature.

**Figure 2.**
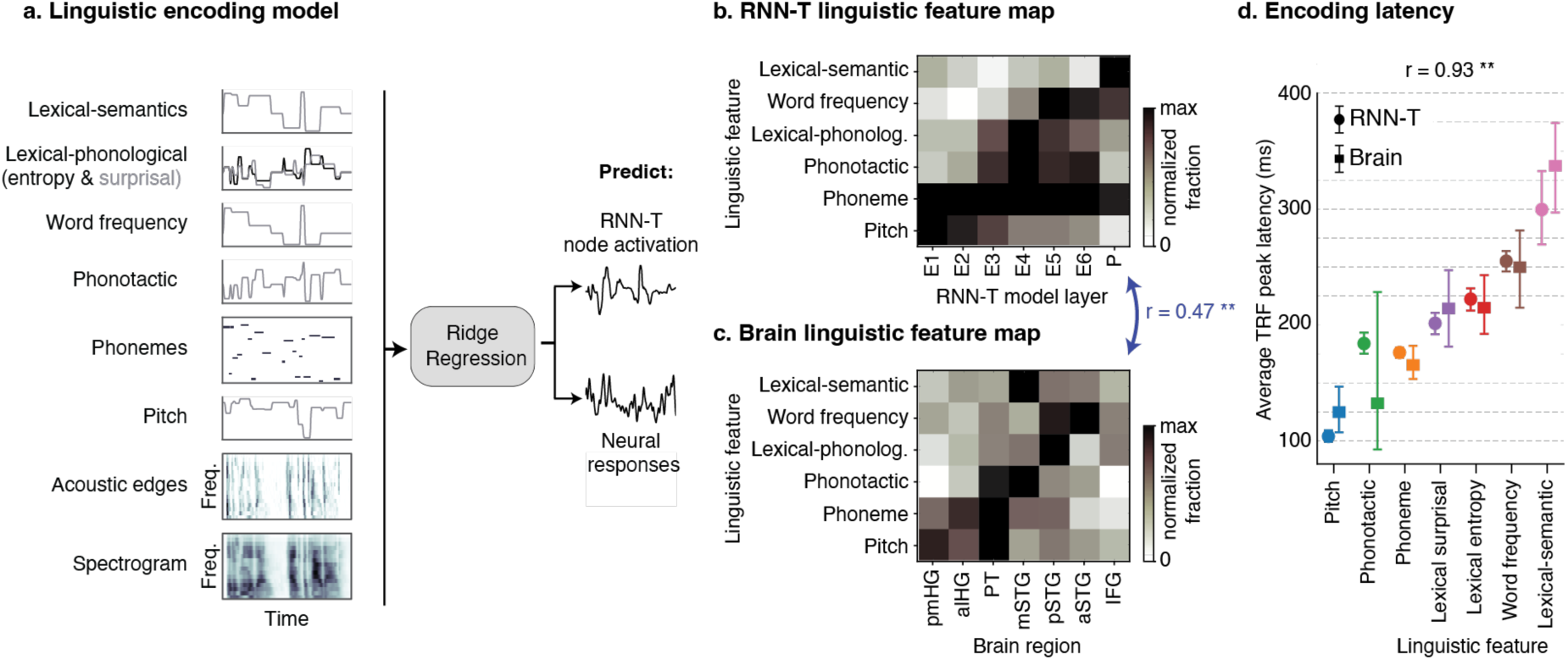
Similar node-level linguistic encoding across the brain and ASR. (a) To measure which linguistic features are encoded in a given electrode site or ASR node, we fit a Ridge regression (TRF) model to predict the response to the stimulus from a set of time-aligned acoustic and linguistic features. This set includes pitch, phonemes, phonotactics, lexical entropy and surprisal, word frequency, and semantic neighbourhood density. The contribution of each feature was assessed by replacing it with a shuffled control and measuring the resulting drop in prediction accuracy. (b) The normalized fraction of nodes in each model layer (E1–E6, P) that significantly encode a given linguistic feature. (c) The normalized fraction of electrodes in each anatomical region that significantly encode a given linguistic feature. The Spearman correlation between the RNN-T linguistic feature map and the brain linguistic feature map is 0.473, p = 0.0016, where ** indicates p < 0.01. (d) The temporal order of linguistic feature encoding in both systems, calculated as the average latency of the peak TRF weight for each feature. Error bars indicate 95% confidence intervals (bootstrap, N=1000). The Pearson correlation between the average encoding latencies of the brain and the RNN-T model is 0.93, p = 0.0027, where ** indicates p < 0.01.

Figure 2b shows the normalized fraction of nodes in each ASR layer that significantly encode each linguistic feature group (fractions divided by maximum across feature/row). Figure 2c shows the same, but for the normalized fraction of neural sites in each brain region. We find a significant correlation between the feature representations over the 7 model layers and the 7 brain regions (Spearman’s 𝑟 = 0.473, 𝑝 = 1.6×10^−3^). To rigorously quantify the alignment of this hierarchical progression, we calculated the center of mass (weighted mean depth) for each linguistic feature within both systems. For the model, this metric defined the average layer (E1–P) where a feature was most strongly encoded; for the brain, it defined the average anatomical region along the ventral stream (pmHG to IFG). We found a strong, significant linear relationship between a feature’s computational depth and its biological depth (Spearman’s r = 0.88, p = 0.033). This result quantitatively confirms that despite their distinct architectures, both the ASR model and the human brain prioritize the extraction of linguistic features in a highly correlated, parallel order, progressing from acoustic to lexical to semantic representations. Supplementary figure S2 shows the average t-statistic without thresholding, normalized by the maximum of each row (feature). We observed that the brain results show broader encoding of higher-level linguistic features in the nonprimary auditory cortex (PT and STG) compared to the primary auditory cortex (HG). This trend is also mirrored in the ASR model. These trends do not depend on the specific choice of the threshold value, as we can see a similar result using average t-values of groups instead of fractions.

Figure 2d shows the temporal order of the encoding of these linguistic features in the ASR model and the brain. To measure the encoding latency for each feature, we first select the group of nodes/electrodes that significantly encoded that feature. For each node/electrode, we compute the latency of the peak absolute value of the TRF weight associated with that feature. We average the latencies to obtain a single latency value for each feature. Finally, using bootstrapping, we obtain 95% confidence intervals for the average latency of each feature. In line with prior findings in the human brain (*30*), the results show a particular temporal order of emergence for the different levels of linguistic information in both systems (Pearson correlation between average latencies of features in the brain and the RNN-T ASR model, r = 0.93, p = 0.0027). To ensure that this temporal order does not depend on the method of latency estimation, we also measure the onset latency from the TRF weights and uncover a nearly identical hierarchical ordering of the linguistic features in both the ASR model and brain (Supplementary Fig. S3).

### Population-level linguistic encoding in the ASR

While the regression approach enabled us to compare the results between the two systems, it cannot capture population-level representations. This is especially relevant in the case of the ASR model, since we have full access to the entire neural population. To determine the patterns of linguistic encoding in the model, we decoded each linguistic feature from the activations of the model (Figure 3a). Because linguistic features are defined at different resolutions and linguistic units have different lengths, we performed a unit-aligned analysis. For example, to predict linguistic features that are defined at phoneme resolution—pitch, phoneme, bi-phone probability, lexical entropy and surprisal—we extracted a layer’s activations at time points {𝑝_𝑖_ + 𝛿}, where {𝑝_𝑖_} were the time points corresponding to phoneme centers and 𝛿 was a constant time lag. Similarly, for word resolution features—word frequency, semantic neighborhood density, contextual embedding—we extracted activations at time points {𝑤_𝑖_ + 𝛿}, where {𝑤_𝑖_} were the time points corresponding to word centers and 𝛿 was a constant time lag. For predicting the categorical phoneme labels we fit linear Ridge classifiers, and for the rest of the features, Ridge regressors (*30*). The best time lag (𝛿) and regularization parameter (𝜆) were determined independently for each layer-feature pair by maximizing the 10-fold cross-validated prediction scores.

**Figure 3.**
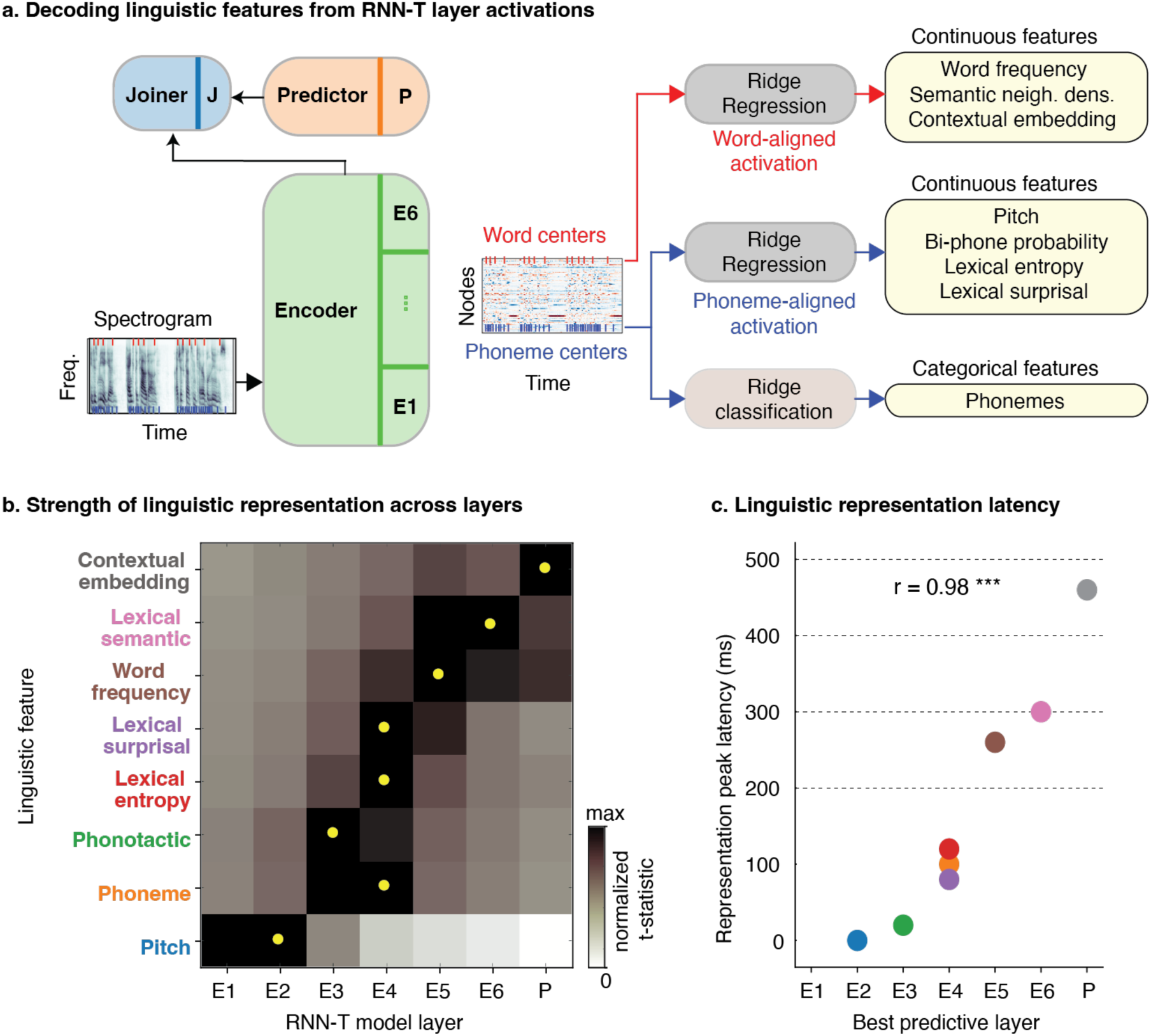
Population-level linguistic encoding in the ASR. (a) To measure how strongly linguistic features are encoded at the population level (layer embedding), we trained single-lag Ridge regressors (for continuous features) or classifiers (for categorical features) to decode specific linguistic features from the activations of each model layer (E1–E6, P). Activations were extracted at time points corresponding to either phoneme centers (blue) or word centers (red), depending on the feature’s temporal resolution. (b) The min-max normalized prediction score for each linguistic feature across model layers (normalized between the baseline spectrogram prediction score and the maximum score across all layers). Yellow dots represent the peak layer for each feature. (c) The emergence of linguistic representations in the model, plotted as the best predictive layer for each feature against its corresponding optimal decoding latency. The Spearman correlation between layer depth and linguistic representation latency is 0.98, 𝑝 = 3.4×10^−5^, where *** indicates p < 0.001).

We also decoded each linguistic feature from the 256-dimensional baseline predictor described earlier. Figure 3b shows the prediction scores of predicting each feature from each layer of the model, normalized per feature such that zero corresponds to the baseline prediction score and 1 corresponds to the best score across all layers of the model. We can see that as we move deeper into the model, representations of higher-order linguistic information emerge. Based on these scores, we can associate each linguistic feature with a layer of the model and find the best lag for that layer-feature pair. As a result, we can describe the place of a linguistic feature in the speech recognition process by its time lag and layer depth (Figure 3c). Each linguistic feature’s best predictive layer depth and the feature’s representation peak latency are strongly correlated (Spearman’s 𝑟 = 0.98, 𝑝 = 3.4×10^−)^). Together, these results show a pronounced emergent linguistic representation in the model through time and space and with a specific order, enabling a direct comparison with the results observed in the brain.

### Effect of model training on linguistic representation

Finally, we explored the conditions in which such a linguistic representation can emerge in the model. We tested two hypotheses: the emergent representation is training-dependent, and the emergent representation is language-dependent. To test these, we decoded the same list of English linguistic features from three different uni-directional RNN-T models: one trained on English, one on French, and a randomly initialized model which underwent no training (Figure 4a). The results show a stark difference between the English and random models, where the random model does not show a strong linguistic representation for any feature compared to the baseline, especially for deeper layers. The French model shows a very similar representation of pitch compared to the English model and a relatively strong representation of only pitch, phonemes, and phonotactics compared to the random model.

**Figure 4.**
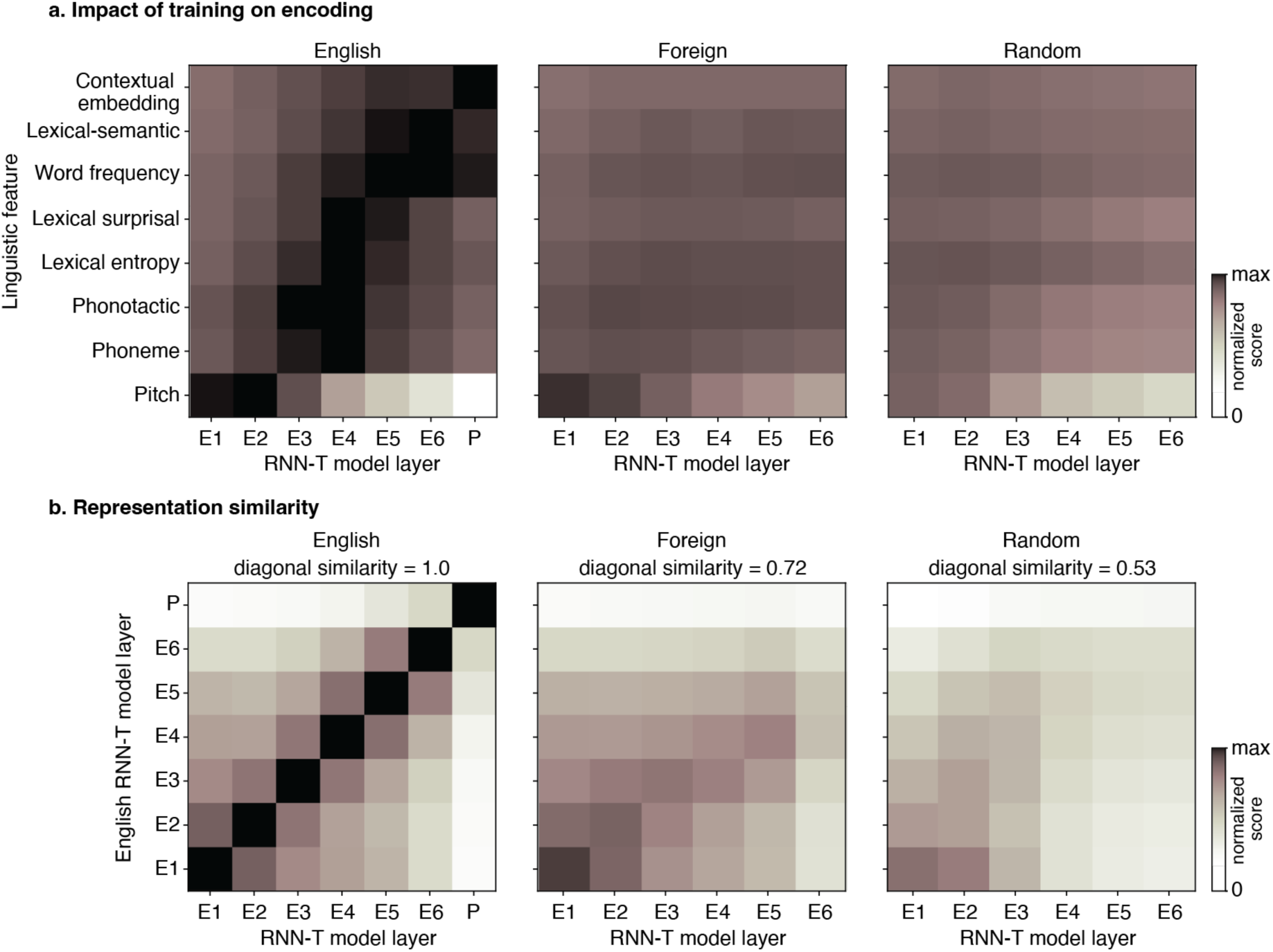
Effect of model training on linguistic representation. (a) Linguistic decoding performance (as in Figure 3) across three uni-directional RNN-T models: one trained on English, one trained on French (“Foreign”), and a randomly initialized model (“Random”). The Predictor branch (P) was excluded for Foreign and Random models because their top-down representations cannot be aligned to the English transcripts. Values are min-max normalized between the baseline (spectrogram) prediction score and the maximum score across all layers of all models. (b) Pairwise centered kernel alignment (CKA) similarity matrices comparing the layer activations of the English RNN-T model (y-axis) against the layer activations of each of the three RNN-T models (x-axis): English, Foreign, and Random. The average CKA similarity along the diagonal is displayed as the diagonal similarity for each model with the English model.

We can also compare these models more directly by computing the similarities of their representations of the same stimulus using centered kernel alignment (CKA) (*31*). We compared the representations of each layer of all three models to the layers of the English model (Figure 4b). We observed that the English and French models share similar representations in their first two layers to a large degree but steadily decrease in similarity over layers overall. In contrast, the random model has weak similarity to the English model, even from early layers. To summarize the similarity of each with the English RNN-T model, we computed the average CKA similarity along the diagonal, finding that this diagonal similarity decreased from 1.0 in the ideal case (English-vs-English) to 0.72 for the French model and 0.53 for the random model. Put together, these results show that the emergent linguistic representation observed in the RNN-T is both training-dependent and language-dependent.

## Discussion

This study reveals a striking convergence in the computational strategies of the human brain and artificial speech recognition systems. By employing a biologically constrained, causal RNN- Transducer, we demonstrate that the model spontaneously develops a representational hierarchy that mirrors the cortical ventral stream, not only in its spatiotemporal organization but, critically, in the specific linguistic content encoded at each stage. Unlike prior works restricted to text-based language models or non-causal architectures (*6–11*), our approach encompasses the full trajectory of speech processing, from acoustic input to semantic understanding. This establishes that the brain’s hierarchical transformation of sound into meaning is likely an efficient computational solution shared by both biological evolution and task-optimized artificial learning, rather than an artifact of specific biological hardware.

### Biological Plausibility and Emergent Hierarchy

Our claim of biological plausibility requires a careful distinction between a model’s physical implementation and its computational strategy. While our model diverges from neurobiology at the implementational level, using backpropagation for learning (*32*, *33*) and possessing a simplified feedback structure (*34*, *35*), it shows compelling parallels at the algorithmic level. The model’s input is a Mel spectrogram, a bio-inspired representation mimicking the cochlear output (*36*, *37*). From this plausible starting point, the model’s end-to-end training allows it to discover its own processing hierarchy, unlike traditional multi-stage ASR systems where such stages were pre-defined.

This hierarchical progression is consistent with patterns identified in prior studies using other architectures (*38–41*), suggesting a robust phenomenon. However, it is important to distinguish this approach from simpler encoding models like Spectro-Temporal Receptive Fields (STRFs) (*29*, *42*). While STRFs offer a biologically realistic account of individual neural sites (*37*, *42*, *43*), they are not designed to perform the overarching computational task of speech recognition. By using a task-optimized RNN-T, we provide a computationally explicit hypothesis for *why* these representations emerge: they are robust solutions for solving the sound-to-meaning problem under causal constraints. Interestingly, we find that low-level acoustic features like pitch strongly emerge in the initial layers (Figure 2b), creating a functional parallel to the early auditory pathway before abstract linguistic processing begins.

### Quantitative Alignment of Spatiotemporal Hierarchies

Our findings move beyond demonstrating a simple correlation between model layers and brain regions. We show not only that model layers map topographically onto the cortical hierarchy, but also that the representational content at each stage is analogous and follows a statistically correlated trajectory (r=0.88) from acoustic to semantic processing. Furthermore, this progression unfolds over a similar timescale (Figure 2d), with deeper model layers mapping to cortical regions that are both anatomically downstream and temporally later in their neural response.

This alignment suggests that the observed hierarchical transformation is a robust solution for processing speech under realistic temporal constraints. While prior work has characterized representational alignment through regression, our identification of specific linguistic features across the hierarchy adds crucial depth. The fact that this distinct artificial system converges on a representational hierarchy so similar to the brain’s, both spatially (Figure 1e) and temporally (Figure 2d), suggests they have arrived at a common, efficient computational strategy.

### Mapping to the Cortical Ventral Stream

Our findings show a striking alignment between the ASR model’s hierarchical layers and the cortical ventral stream, the primary pathway for speech comprehension (*44–46*). This alignment is quantitatively supported by the monotonic increase in both neural latency and anatomical distance from the primary auditory cortex as a function of model depth. The progression from early to deep model layers mapping from posterior to anterior temporal and inferior frontal regions aligns well with the ventral stream’s role in the sound-to-meaning transformation.

We also observed correlations between deeper model layers and regions of the dorsal stream (*44–46*), though the interpretation is less direct given the dorsal stream’s role in speech production. This may reflect shared computations for complex sequential information (*47–50*) or engagement of verbal working memory. Additionally, the Inferior Frontal Gyrus (IFG) most strongly matched the later layers and the predictor branch of the ASR model. Given the IFG’s implicated role in unifying lexical and syntactic information (*47–49*, *51*, *52*), this match suggests that the model’s predictor branch functionally mirrors the brain’s high-level integration mechanisms. While the sentence repetition component of our task may have influenced representations in IFG through its role in verbal working memory and cognitive control (*53–56*), this confounding effect should be small since only a small fraction of sentences were repeated. Overall, our findings indicate a functional congruence between the RNN-T ASR model and the human brain’s hierarchical linguistic representations (*44*, *57–59*), advancing prior work by highlighting the specific regions and representational latencies at each stage of this hierarchy which are common across both the human brain and the ASR model.

### The Role of Training and Language Specificity

Our analysis of control models yielded new insights into how this hierarchy forms. We found that in a French-trained model, acoustic features like pitch were strongly represented for English stimuli, but higher-level linguistic representations were weak, and they were absent in an untrained random model. This demonstrates that the high-level linguistic hierarchy is not an inevitable byproduct of deep architectures and longer temporal integration (*60*) but is critically dependent on language-specific training.

Our comparison of layer representations revealed that the French model diverged from the English model quickly after the first two layers, pinpointing where language-specific encoding begins. These results support recent findings that models trained on native languages better predict neural responses (*6*) and offer a granular explanation: the divergence is driven by the failure of non-native models to develop higher-order lexical and semantic features for the target language.

### Limitations and Future Directions

A key distinction is that the RNN-T processes speech in a non-lateralized fashion, whereas the brain shows a strong left-hemisphere bias for higher-level linguistic tasks (*27*, *28*). Our single-pathway model more closely approximates the computational style of the left hemisphere; the weaker correspondence in the right hemisphere may reflect the model’s inability to capture the distinct, parallel processing strategies and speech comprehension capabilities of the right hemisphere (*61–65*). Future biologically accurate models may require dual-pathway architectures or multitask objectives (*38*) to capture these complex hemispheric dynamics.

Additionally, ASR models trained with a recognition loss tend to become speaker-independent, likely removing speaker-specific information such as pitch in deeper layers (*66*, *67*). This matches our finding that pitch encoding fades in later layers. Despite this, complex semantic representations emerge from the relatively simple objective of predicting graphemes, a hallmark of deep learning systems. Future work could systematically investigate how the choice of the model’s output unit (e.g., graphemes, phonemes, or words) modulates the specific nature and strength of these emergent representations, but our results establish that the core brain-like hierarchy is a robust outcome of the end-to-end learning process.

In conclusion, this study bridges neuroscience and artificial intelligence by revealing how both systems process speech in hierarchical layers. Our findings not only advance our understanding of the brain’s mechanisms for linguistic encoding but also offer valuable guidance for the development of more sophisticated and biologically informed ASR models. Moving forward, integrating more brain-like features into artificial systems may unlock new possibilities for understanding and enhancing human and machine communication.

## Methods

### Participants, neural data, task, and stimuli

Fifteen human patients (7 female, mean age: 36, range: 19-58) with drug-resistant epilepsy were studied. All patients were implanted with intracranial electroencephalography (iEEG) electrodes for epileptogenic foci localization. Twelve of the patients had stereoelectroencephalographic (sEEG) depth electrodes, while the other three had both depth electrodes and subdural grids and/or strips. All recordings were inspected by an epileptologist to ensure they were free of interictal spikes. The patients provided written, informed consent to participate in the research study prior to implantation, and the protocol was approved by the institutional review board at the Feinstein Institutes for Medical Research.

The subjects listened to approximately 30 minutes of stories spoken by voice actors. Occasional pauses in the story were added, resulting in 53 trial segments, and the subjects were asked to repeat the most recent sentence before the pause to ensure they were paying attention. All subjects were able to repeat the sentences without issue. iEEG signals were acquired at 3 kHz sampling rate, and the envelope of the high-gamma band (70-150 Hz) was extracted with the Hilbert transform (*68*). This signal was then z-scored and resampled to 100 Hz.

### Electrode localization

Electrodes were localized using the iELVis toolbox (*69*). Prior to the iEEG recordings, each patient underwent a T1-weighted 1 mm isometric structural magnetic resonance imaging (MRI) scan on a 3 T scanner. After the electrode implantation, a computed tomography (CT) scan together with a T1-weighted MRI scan at 1.5 T were acquired. The post-implantation CT and MRI scans were co-registered to the preoperative MRI scan using FSL’s BET and FLIRT algorithms (*70–72*). Afterwards, the artefacts of the contacts on the co-registered CT were identified manually in BioImageSuite (*73*). Volumetric information was obtained by processing and reconstructing the T1 scan using FreeSurfer v.6.0 (recon-all command) (*16*). For the anatomical analyses across participants, we mapped the coordinates of the electrodes for each participant to the FreeSurfer average brain (fsaverage), which is a template brain based on a combination of MRI scans of 40 real brains.

### Electrode selection

Electrodes were projected onto the nearest cortical surface and assigned to a subregion according to the Destrieux cortical atlas (*16*, *74*). We chose the Destrieux atlas due to its finer granularity in the regions of interest. We selected all electrode sites within the auditory cortex (AC; 𝑁 = 335) —specifically, HG, which includes the PAC or core (*75*); the PT, which coincides with the parabelt; and the STG—and the inferior frontal gyrus (IFG; 𝑁 = 191). For increased granularity during analysis, we further split the HG into posteromedial (pmHG) and anterolateral (alHG) sections, as they have been shown to exhibit different properties (*76*). For the same reason we also split the STG into three sections of anterior (aSTG), middle (mSTG), and posterior (pSTG). The border between mSTG and pSTG was defined as the crossing of a virtual line extending from the transverse temporal sulcus with the lateral surface of the STG (*76*, *77*). The border between mSTG and aSTG was set such that they are roughly equal in size. Figure 1a shows the general location of IFG and the subregions of the auditory cortex on the FreeSurfer average brain (*16*).

To determine whether an electrode site was speech-responsive, we first selected 53 pre-trial silence segments ([-1 s, 0 s] relative to segment onset) and using a 200 ms wide non-overlapping moving average window reduced it into 53 × 5 data points. We then selected 53 post-onset speech segments ([0.4 s, 1.2 s] relative to segment onset) and using a similar moving average window reduced it into 53 × 4 data points. We performed a two-sample t-test between the two distributions for each electrode to obtain a p-value, and then performed a Benjamini-Hochberg (false discovery rate) correction with an alpha of 0.05 to determine the speech-responsive electrodes. This method was observed to be more robust compared to some alternatives when viewing across the entire regions of interest which included electrodes inside and outside the auditory cortex with different response latencies.

We limited our analysis to sites that were determined to be speech-responsive (𝑁 = 291/526). We labeled the neural sites in both hemispheres based on their anatomical region to enable population tests: posteromedial Heschl’s gyrus (pmHG; 𝑁_𝐿_ = 12/15, 𝑁_𝑅_ = 16/19), anterolateral Heschl’s gyrus (alHG; 𝑁_𝐿_ = 34/36, 𝑁_𝑅_ = 32/34), planum temporale (PT; 𝑁_𝐿_ = 9/12, 𝑁_𝑅_ = 18/20), middle superior temporal gyrus (mSTG; 𝑁_𝐿_ = 44/53, 𝑁_𝑅_ = 17/18), posterior superior temporal gyrus (pSTG; 𝑁_𝐿_ = 27/32, 𝑁_𝑅_ = 21/23), anterior superior temporal gyrus (aSTG; 𝑁_𝐿_ = 19/60, 𝑁_𝑅_ = 0/13), and inferior frontal gyrus (IFG; 𝑁_𝐿_ = 32/119, 𝑁_𝑅_ = 10/72). The electrode locations and their responsiveness are plotted in Figure 1b on the average FreeSurfer brain, where the color indicates whether an electrode is speech-responsive.

### RNN-Transducer

To model the speech processing mechanism, we use an RNN-Transducer (RNN-T) (*13*), a recurrent neural network trained to perform automatic speech recognition, i.e., predicting graphemes from the speech spectrogram. The RNN-T is a general architecture, which can also utilize self-attention and transformer layers, but our instantiation uses causal LSTM layers. The model consists of two branches—the encoder and the predictor (Figure 1c). The encoder branch, consisting of six uni-directional LSTM layers of 640 nodes each (E1–E6), acts as a bottom-up encoder. It transforms the input speech spectrogram into a representation used to predict the output labels. The predictor branch, consisting of a single uni-directional LSTM layer of 1024 nodes (P), acts as an internal language model. It stores an internal representation of the prior predictions of the model that, together with the encoder’s output, are used to make the next prediction. A joint network merges the final encoder branch layer with the predictor branch’s embedding and predicts text characters to generate the speech transcription. The English network outputs are projected to 42 logits, corresponding to 41 characters plus BLANK. Similarly, for French, we use 72 output units. The model is trained with the RNN-T loss on the Switchboard and Fisher datasets, which, put together, consist of about 2,000 hours of English phone conversations. As presented in prior work (*78*), an additional 4-fold data augmentation was applied to the input spectrograms to allow better generalization of the model, bringing the duration of the unique training data to 10,000 hours. These augmentations included speed and tempo perturbations which changed the rate of speech in the interval [0.9, 1.1] with or without altering the pitch or timbre of the speaker, as well as spectrogram augmentation which masked the spectrum of a training utterance with a random number of blocks of random size in both time and frequency (*78*). Augmentation with an external language model as discussed in (*78*) was not employed here. We computed layer activations for all layers of the model in response to the same 30-minute stimulus set that the human participants listened to.

### Time-aligning RNN-Transducer activations to neural data

The processed neural data is sampled at 100 Hz and has shape 𝑇 × 𝐶, where 𝑇 is the number of time steps and 𝐶 is the number of electrode channels. The activations of the layers in the encoder branch of the RNN-T models are sampled at 50 Hz and have the shape 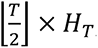, where 𝐻_𝑇_ is the number of nodes in each hidden encoder layer. These activation matrices can be trivially aligned with the neural data sampled through a resample with the integer factor 2. The activations of the predictor branch, however, have shape 𝑈 × 𝐻_𝑃_, where 𝑈 is the number of output graphemes and 𝐻_𝑃_ is the number of nodes in the hidden prediction layer. These activation matrices cannot be directly aligned with the neural data since they are based on graphemes instead of time. To make this alignment possible, we need to find the most likely “time warping” between output grapheme indices (1 ≤ 𝑢 ≤ 𝑈) and time 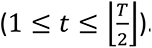. We run the forward-backward algorithm defined in section 2.4 of Graves (*13*) on the grapheme output probability lattice produced by the model to obtain this “time warping” between the two sequences. We then use the 𝛼: 𝑡 → 𝑢 alignment to stretch the activation matrix 𝑍_𝑃_ with shape 𝑈 × 𝐻_𝑃_ into a matrix 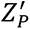 with shape 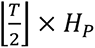 where 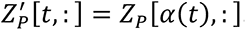. Since 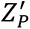 has the same shape as encoder layer activations, we can trivially align it to the neural data.

### Linguistic features of speech

#### Pitch

As a measure of pitch, we computed the pitch contour (F0) of the speech signal using the PyWORLD python package, which is a python wrapper for the WORLD vocoder (*79*). We then averaged the value across the duration of each phoneme to obtain a phoneme-average pitch.

#### Phonemes

For phonetic features, we used the categorical (one-hot encoded) representation of ARPAbet phonemes. We chose this because it allows classification in the layer-level analysis.

#### Phonotactic features

Phonotactics represent phoneme transition probabilities, so we used the biphone probability 𝑃_𝑎𝑏_ for phoneme bigram 𝑎𝑏:

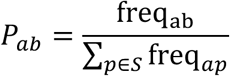

where 𝑆 is the set of all ARPAbet phonemes. To compute the frequencies of each biphone, we used the CMU dictionary (http://www.speech.cs.cmu.edu/cgi-bin/cmudict) to convert words to phoneme sequences and counted the occurrence of each biphone using the SUBTLEX-US corpus, which is an English word frequency dataset calculated from movie subtitles (*80*). Since biphone frequencies were calculated from a word frequency dataset and without access to word transition probability information, we counted the first phoneme transition of words separately from non-first phonemes. For example, the biphones for the phrase “red hat” are the following: /#r/, /re/, /ed/, /#h/ (not /dh/), /hæ/, and /æt/. The frequency of a phoneme bigram represents the degree of exposure of an average native listener to the bigram and measures its probability in natural speech. We purposefully chose a non-position-specific measure of phonotactics (as opposed to the more common approach (*81*)) to maximally dissociate this effect from lexical processes. This level represents the expectation and surprisal of the listener when hearing a new phoneme, based on the immediate past. This prelexical phonotactics feature could indicate predictive coding mechanisms that operate on the phonemic level (*82–85*).

#### Lexical-phonological features

To measure the lexical-phonological effect, we used lexical entropy and surprisal. These values were calculated for each phoneme within a word from the previous phonemes in that word. The surprisal caused by phoneme φ_i_, S(𝑖), in word w=φ_1_…φ_K_ indicates the improbability of hearing phoneme φ_i_ based on the previous i−1 phonemes that came before it in the word and is calculated as follows:

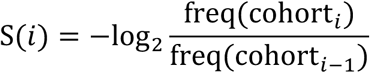

where freq(cohort_𝑖_) is the summed frequency of all words that start with the phoneme sequence φ_1_…φ_i_. On the other hand, the lexical entropy, E(i), for phoneme φ_i_, is the entropy within all words that start with the phoneme sequence φ_1_…φ_i_ (the cohort) (*86*):

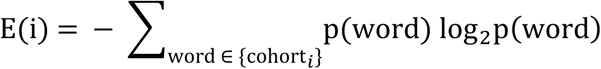

where p(word) indicates the relative frequency of the word within the cohort. These two parameters together encode the incremental lexical competition among all phonologically consistent candidates as a word is being heard, weighted by their frequency. To compute lexical surprisal for the word-initial phoneme, we assumed a transition from the entire lexicon, i.e., how surprising it is to hear a word starting with phoneme p given the frequencies of all the words in the lexicon.

#### Lexical-semantic features

To study the encoding of semantic information, we represented each word with its semantic neighborhood density (SND) obtained from the English Lexicon Project (*87*, *88*), which refers to the relative distance between a word and its closest neighbors based on a global co-occurrence model (*87*, *88*). The neighborhood density can encode the degree of activation of semantically related words in the lexicon upon hearing the target word.

#### Contextual embedding

We used the embedding obtained from the last hidden layer of a pre-trained GPT-2 XL model obtained from Hugging Face (*89*). This 1.5B parameter version of GPT-2, a transformer-based language model was pretrained using a causal language modeling (CLM) objective on English language data. CLM is a training goal where the model predicts the next token in a sequence given its preceding tokens, ensuring that the prediction for a position can only depend on known outputs at previous positions.

To associate contextual embedding to words in the analysis data, we concatenated the transcripts for all 53 trials, in the same order the participants heard them. Then, for each word, we gave the model the last token of the target word and the 511 tokens preceding it (if available) to obtain a contextual embedding for that word. The total 512 tokens are half the maximum context window size of GPT-2 XL and correspond to about 3 minutes of context in the experiment data.

#### Predicting neural responses from ASR activations

We predict the neural activity recorded at each electrode from each of the layers of the ASR model (Figure 1c). We map ASR layers instead of ASR nodes to electrodes, because the high-gamma envelope of an iEEG electrode represents a readout of the activity of neighboring neural populations consisting of thousands of neurons rather than individual neurons.

To predict a neural response from a layer of the model, we fit a linear single-lag Ridge regressor that predicts the electrode activity from the layer activations in response to the same stimuli. Discounting a constant lag term 𝛿, the output at time 𝑡 is predicted only from the input at time 𝑡 + 𝛿. In other words, 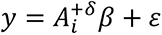, where 𝑦 is the 𝑇 × 1 neural response vector, 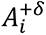 is the 𝑇 × 𝐻 activation matrix for layer 𝑖 of the ASR model with 𝐻 nodes shifted by 𝛿 time steps across its first dimension, 𝛽 is the 𝐻 × 1 regression model that maps 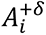 to 𝑦, and 𝜀 is the regression error.

We use the Ridge regressor from the python scikit-learn package to fit these models. The optimal lag value (𝛿) and ridge regularization parameter (𝜆) are chosen independently for each electrode- layer pair by maximizing the prediction score through 10-fold cross-validation. Here, and in all regression models in this paper, nested cross-validation was used to fit models in an unbiased way by performing finding the optimal lag and regularization parameters within only the training set.

#### Predicting neural responses and ASR activations from linguistic features

We used a temporal receptive field (TRF) (*29*) framework to measure the extent to which different linguistic features are encoded in the neural responses recorded from the brain and activations extracted from the ASR nodes (Figure 2). We used a broad spectrum of acoustic and linguistic features as predictors: spectrogram, acoustic edges, pitch (phoneme-average F0), phonemes, phonotactics (− log 𝑃_𝑎𝑏_), word frequency, lexical-phonological (lexical entropy and lexical surprisal), and lexical-semantics (semantic neighborhood density). The enormous dimensionality of the contextual embedding feature prevented its use in this framework.

To produce a distribution of shuffled models for statistics, we replaced each linguistic feature with a shuffled version, one at a time, 100 times. We measured a t-statistic of encoding using a one-sample t-test to compute a t-value of the difference in scores between the true features and the shuffled versions. A negative value for the t-statistic is possible if the shuffling produces a random feature that happens to be better encoded in the neural response than the true stimuli due to the small duration of the stimuli. So, following our previous work with the same data (*30*), we used 𝑡 = 19 as the threshold of significant encoding, since none of the negative t-values were less than 𝑡 = −19. This method was chosen over a non-parametric test of the shuffled score improvements due to the computational limitations of estimating enough cross-validated models to build a robust distribution with good resolution at low p-values. Special care was taken for each type of feature to ensure that the shuffling did not alter the influence of lower-level features in the hierarchy, as described below.

#### Pitch features

We grouped all the words within our 30-minute data based on their number of phonemes. Then we shuffled the word-to-pitch sequence association map within each group.

#### Phonetic features

We took the CMU pronunciation dictionary (http://www.speech.cs.cmu.edu/cgi-bin/cmudict), grouped words by their length measured in phonemes, and then shuffled the word-to-phoneme mapping within each group. As a result, each word will have a consistent pronunciation at every occurrence, but words that share phonemes will have independent pronunciations, e.g. /kæt/ and /bæt/ no longer share two of their three phonemes. We constrained the reassociation to words of same length so that we kept the phoneme alignment information intact and because words of the same length are more similar in frequency of occurrence (i.e. shorter words tend to be more frequent). This is a rather strict control, since shuffling pronunciations with other actual English words maintains the proper syllabic structure for English words.

#### Phonotactic features

To generate controls for phonotactic features, we shuffled the bigram-to-frequency associations (i.e. the look-up table for bigram frequencies), which means that each bigram was associated with the frequency of a randomly chosen bigram from the true distribution. This control scheme maintained consistency across multiple occurrences of the same bigram. To counter the effect of the separation caused by the first vs. non-first phoneme grouping, we perform the above shuffling separately for first phones (ones starting with #) and non-first biphones, so that any first vs. non-first effect will be maintained in the control, and thus discounted.

#### Word frequency

We grouped words based on their phoneme length and shuffled the word-to-frequency associations within each group.

#### Lexical-phonological features

We grouped all cohorts based on the length of their shared phoneme sequence and shuffled the cohort-to-frequency associations within each group. We used this constrained shuffling to keep the effect of secondary information, such as the phoneme position in the word and word length, unchanged. This control scheme also satisfies consistency, i.e. if two words share their first k phonemes, the cohort information for their first k positions would be the same because the same cohorts are mapped to the same information.

#### Lexical-semantic features

The control for the semantic condition was constructed by grouping words based on their phoneme length and shuffling the word-to-SND associations within each group.

#### Predicting linguistic features from ASR activations

We predicted different linguistic features from each of the layers of the ASR model. Because linguistic features are defined at different resolutions and linguistic units have different lengths, we performed a unit-aligned analysis. For example, to predict linguistic features that are defined at phoneme resolution—pitch, phoneme, biphone probability, lexical entropy and surprisal—we extracted a layer’s activations at time points {𝑝_𝑖_ + 𝛿}, where {𝑝_𝑖_} were the time points corresponding to phoneme centers and 𝛿 was a constant time lag. This 𝑁 × 𝐻 activation matrix, where 𝑁 is the number of phonemes and 𝐻 is the number of nodes, was multiplied by an 𝐻 × 𝐷 linear decoder to predict the 𝐷-dimensional linguistic feature. Similarly, for word resolution features—word frequency, semantic neighborhood density, contextual embedding—we extracted activations at time points {𝑤_𝑖_ + 𝛿}, where {𝑤_𝑖_} were the time times corresponding to word centers and 𝛿 was a constant time lag.

For predicting the categorical phoneme labels we fit linear Ridge classifiers, and for the rest of the features Ridge regressors. The best time lag (𝛿) and regularization parameter (𝜆) was determined independently for each layer-feature pair by maximizing the 10-fold cross-validated prediction scores.

#### Representation similarity of ASR model layers

We use two methods to compare representations of two ASR layers. The first is to compare their linguistic decoding results. The second is to compare the representations directly with the centered kernel alignment (CKA) method, which is used to measure the similarity between two sets of high-dimensional vectors, especially representations learned by neural networks (*31*). For this comparative analysis, we used three models: the baseline model trained on English data; a “random” model that has the same architecture as the English model, but randomly initialized and untrained; a “foreign” model that was trained on a French ASR task and was not exposed to English.

## Data availability

Although the iEEG recordings used in this study cannot be made publicly available due to patient privacy restrictions, they can be requested from the author (N.M.).

## Code availability

Code for preprocessing neural data, selecting responsive electrodes, and creating brain plots is available in the naplib-python package (*S0*). A reproducible capsule can be found on CodeOcean (*S1*). The software contains tutorial notebooks and guides for preprocessing data as was done here.

## Acknowledgements

This study was funded by the National Institute on Deafness and Other Communication Disorders, (R01DC014279 to N.M.). S.B. was also supported by the National Institute on Deafness and Other Communication Disorders, R01DC019979. The funders had no role in the study design, data collection and analysis, decision to publish and manuscript preparation.

## Author contributions

Conceptualization (M.K., N.M.); methodology (M.K., G.M., S.T., B.K., N.M.); data collection (S.B., A.D.M.); data analysis (M.K., G.M., S.T., N.M.); writing – original draft (M.K., N.M.); writing – revision (G.M., S.T., B.K., N.M.).

