## Supplementary Fig. S1 for "Parallel hierarchical encoding of linguistic representations in the human auditory cortex and recurrent automatic speech recognition systems"

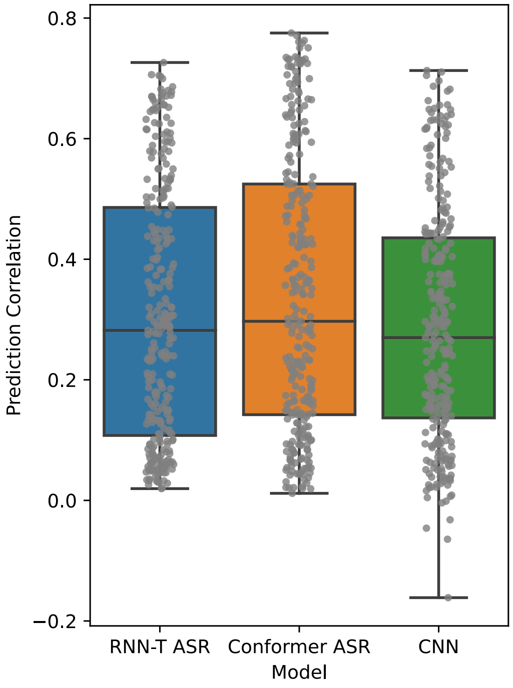

**Figure S1. Prediction correlation of RNN-T ASR model vs Conformer ASR model and purpose-built CNN encoder.**

A 6-layer Conformer-based ASR model was trained using the same training data as the RNN-T, and representations were extracted from its layers for the same stimuli as was given to the RNN-T and human subjects. Ridge regression models were then fit from these representations to predict the neural data using the same cross-validation approach as the RNN-T. The peak prediction scores (across layers) for each electrode are shown in the box-plot distributions for the RNN-T and Conformer models. A CNN encoder model (with nearly identical architecture as Keshishian et al., 2020) was trained to predict neural responses to all speech-responsive electrodes ($N = 291$) from stimulus spectrogram input. The cross-validated prediction correlations are shown in comparison to the scores from predicting the neural data from the ASR model, showing nearly identical performance. The box bounds define the interquartile range, with the center line defining the median, while whiskers indicate the minima and maxima.

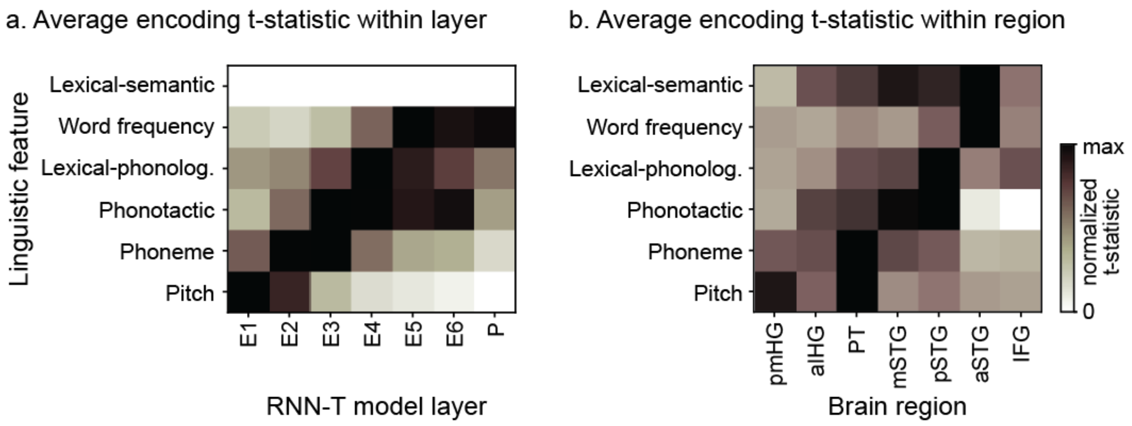

**Figure S2. Average t-values of node-level linguistic encoding in the brain and ASR model.** **(A)** Average t-statistic from nodes within each layer of the RNN-T ASR model. Each row is normalized by the maximum average t-statistic over layers for that row/feature. The mean lexical-semantic t-statistics are negative when averaging across all nodes in each layer of the RNN-T model, potentially because the RNN-T may not tend to represent these high-level features at the individual node level, which leads to the top row appearing entirely 0 after normalization. **(B)** Average t-statistic from electrodes within each brain region. Each row is normalized in the same way as for the RNN-T ASR model.

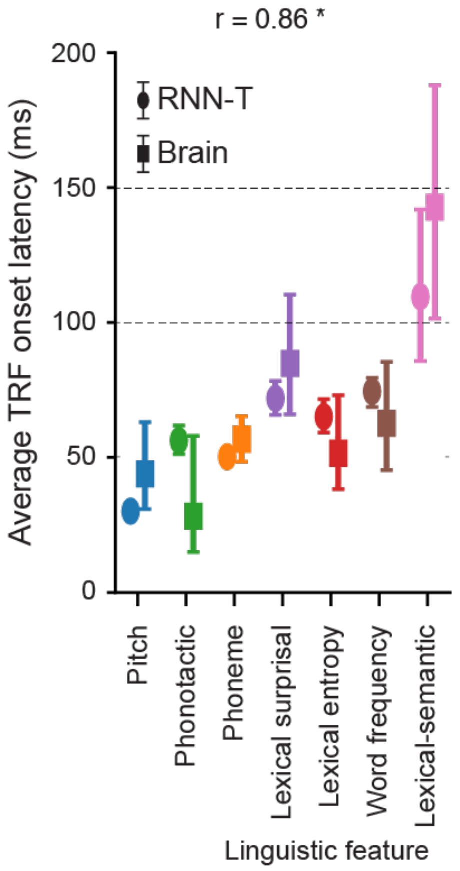

**Figure S3. Latency of linguistic feature encoding in the brain and RNN-T ASR model using a second metric.**

To ensure that the estimated latencies of linguistic features as shown in Figure 2D are not caused by differences in sustained vs temporally sharp responses to certain linguistic features, this figure reproduces that analysis with a new metric for latency. Here, the onset latency of each feature is measured from its TRF weights. For each electrode's TRF for a given feature, the mean and standard deviation of the TRF weights during the 50 ms prior to stimulus onset are computed, with the mean acting as a baseline. Then, the onset latency is determined to be the first time point where the TRF weights surpass the baseline plus twice the standard deviation. A nearly identical hierarchy of feature encoding latency emerges in both the brain and the RNN-T ASR model as with the previous method of using the peak absolute value of the TRF weights. Error bars indicate 95% confidence intervals obtained using bias-corrected and accelerated bootstrap (BCa; $N = 1000$) on the mean latency of encoding across nodes/electrodes. The Pearson correlation between latencies of features in the brain and the RNN-T model is 0.86, p = 0.014, where * indicates p < 0.05.
